# Catecholamine modulation of evidence accumulation during perceptual decision formation; a randomised trial

**DOI:** 10.1101/470120

**Authors:** Gerard M. Loughnane, Méadhbh B. Brosnan, Jessica J.M. Barnes, Angela Dean, L. Sanjay Nandam, Redmond G. O’Connell, Mark A. Bellgrove

**Affiliations:** Trinity College Institute of Neuroscience and School of Psychology, University of Dublin, Trinity College, Dublin 2, Ireland; School of Psychological Sciences and Monash Institute for Cognitive and Clinical Neurosciences (MICCN), Monash University, Melbourne, Australia; Queensland Brain Institute, University of Queensland, Brisbane, Australia

## Abstract

Recent behavioural modelling and pupillometry studies suggest that neuromodulatory arousal systems play a role in regulating decision formation but neurophysiological support for these observations is lacking. We employed a randomised, double-blinded, placebo-controlled, crossover design to probe the impact of pharmacological enhancement of catecholamine levels on perceptual decision making. Catecholamine levels were manipulated using the clinically relevant drugs methylphenidate (MPH) and atomoxetine (ATM) and their effects were compared to those of citalopram (CIT), and placebo (PLA). Participants performed a classic EEG oddball paradigm which elicits the P3b, a centro-parietal potential that has been shown to trace evidence accumulation, under each of the four drug conditions. We found that MPH and ATM administration shortened RTs to the oddball targets. The neural basis of this behavioural effect was an earlier P3b peak latency, driven specifically by an increase in its build-up rate without any change in its time of onset or peak amplitude. This study provides neurophysiological evidence for the catecholaminergic enhancement of a discrete aspect of human decision making, i.e. evidence accumulation. Our results also support theoretical accounts suggesting that catecholamines may enhance cognition via increases in neural gain.

## Introduction

An important line of research in neuroscience aims at understanding the brain mechanisms allowing us to make reliable perceptual judgements based on noisy sensory information. Convergent data from psychophysics, computational modelling and neurophysiology highlight the central role of ‘decision variable’ signals in determining the timing and accuracy of perceptual reports by accumulating sensory evidence over time up to an action-triggering threshold (Gold & Shadlen, 2007; Kelly & O’Connell, 2015; Smith & Ratcliff, 2004; Smith & Vickers, 1988). Such signals have been identified both in the spiking activity of neuronal subpopulations within several sensorimotor regions of the monkey brain (L. Ding & Gold, 2010; Hanes & Schall, 1996) and in signals recorded non-invasively from the human brain (de Lange, Rahnev, Donner, & Lau, 2013; Donner, Siegel, Fries, & Engel, 2009; Kelly & O’Connell, 2013; Loughnane et al., 2016; Newman, Loughnane, Kelly, O’Connell, & Bellgrove, 2017; O’Connell, Dockree, & Kelly, 2012; Philiastides, Heekeren, & Sajda, 2014).

The biophysical mechanisms supporting these accumulation-to-bound dynamics remain to be fully characterised and there is growing interest in the role of neuromodulatory arousal systems (Aston-Jones & Cohen, 2005; de Gee et al., 2017; Murphy, Robertson, Balsters, & O’Connell, 2011; Nieuwenhuis, Aston-Jones, bulletin, 2005, n.d.). Specifically, psychopharmacological studies in which catecholaminergic (i.e., dopamine and noradrenaline) function has been modulated demonstrate improved behaviour on perceptual tasks (Brumaghim & Klorman, 1998; Linssen, Sambeth, Vuurman, & Riedel, 2014; Naylor, Halliday, Psychopharmacology, 1985, n.d.), possibly mediated by an increase in neural gain (Eldar, Cohen, & Niv, 2013; Warren et al., 2016).

However, our understanding of the parameters of perceptual decision-making impacted by catecholamine modulation remains limited. Pharmacological enhancement in the speed of response to stimuli could arise from several sources according to the computational modelling of behaviour, such as the onset or build-up rate of evidence accumulation, or a lowered decision bound (Fosco, White, & Hawk, 2016; Ratcliff & McKoon, 2008). Here we sought to use neurophysiology to adjudicate between these model predictions.

We traced decision formation via the P3b component of the human event-related potential. In recent studies we demonstrated that the P3b exhibits the key characteristics of a build-to-threshold, evidence accumulation process. Like the decision signals that have been observed in intracranial recordings, the P3b builds gradually during decision formation, at a rate that scales with the difficulty of the perceptual decision and reaches a stereotyped amplitude immediately prior to response execution (Kelly & O’Connell, 2013; O’Connell et al., 2012). These dynamics have been observed during extended motion discrimination and contrast change detection judgments but are also apparent during performance of the classic oddball task (Twomey, Murphy, Kelly, & O’Connell, 2015).

We probed catecholamine function using methylphenidate (MPH) and atomoxetine (ATM). MPH and ATM increase monoamine levels to varying extents in different brain regions. In prefrontal cortex, the action of both MPH and ATM converge to increase levels of both dopamine and noradrenaline, initially via reuptake inhibition of the noradrenaline transporter (NET) (Berridge et al., 2006; Bymaster, 2002; Han & Gu, 2006). Within subcortical regions such as the striatum, however, their action diverges with only MPH inhibiting reuptake via the dopamine transporter (DAT) (Volkow, Fowler, Wang, Ding, & Gatley, 2017; Volkow et al., 2001).

We employed a randomised, double-blinded, placebo-controlled, crossover design, manipulating monoamine levels using the clinically relevant drugs MPH, ATM and the serotonin reuptake inhibitor citalopram (CIT), in comparison to placebo (PLA). Citalopram acted here as an active pharmacological control to establish the specificity of any catecholamine modulation.

Participants performed an EEG-oddball paradigm to elicit a neural correlate of this evidence accumulation process, the P3b, under each of the four drug conditions. We hypothesised firstly that MPH and ATM would improve reaction times relative to placebo, whereas no effect of CIT would be observed. Secondly, we hypothesised that enhanced perceptual decision making with MPH and ATM would be underpinned by a decrease in the P3b peak latency, driven via an increase in the rate of evidence accumulation. Thirdly, we examined the Visual Evoked Potential to discern the temporal locus of any catecholaminergic effects on visual processing. Given the lack of evidence in past work for effects on VEP components such as P1 and N1 (Linssen et al., 2014), we hypothesised that these components would not be modulated by our pharmacological manipulations.

## Materials and Methods

### Participants

Forty non-clinical volunteers completed a larger study investigating the influence of monoamine reuptake inhibitors on executive function (Barnes, O’Connell, Nandam, Dean, 2014, n.d.; Dockree et al., 2017). Participants were recruited via advertisements at The University of Queensland, Queensland, Australia. All participants were male, Caucasian, right-handed, had normal or corrected-to-normal vision, aged 18-45 years, and were prescreened by a consultant psychiatrist (L.S.N) for suitability. Individuals were excluded from participation if they reported any history of psychiatric or neurological illness (including head injury resulting in unconsciousness), previous or current use of psychotropic medication, current smoking or use of nicotine products, or significant drug use (significant was defined here as: (i) use of any illicit substances within the last month; (ii) >5 lifetime intake of any illicit drug except cannabis; or (iii) more than monthly cannabis intake, alcohol dependence (>24 units/week)). Before commencing their first session, all participants were screened by a consultant psychiatrist (L.S.N.). The psychiatrist also administered the M.I.N.I. screen (Sheehan et al., 1998). Seven participants were excluded from the analysis due to illness (n=4), technical data acquisition issues (n=1), and behavioural outlier criteria (n=2), during one session. All participants were recruited according to the principles of the Declaration of Helsinki and in accordance with the ethical guidelines of The University of Queensland.

### Drug administration

Each participant attended four sessions, each at the same time of the day, spaced at least 1 week apart. At each session, a single blue gelatine capsule containing methylphenidate (30 mg, mixed dopaminergic and noradrenergic action), atomoxetine (60 mg, mixed dopaminergic and noradrenergic action), citalopram (30 mg, primarily serotonergic action) or placebo (dextrose) was administered. Cognitive testing began 90 min following drug administration coinciding with peak plasma concentrations of the drugs (Hennig & Netter, 2002; Müller et al., 2005; Sauer, Ring, & Witcher, 2005). Subjective side effects of the drug were measured using a visual analogue scale (VAS) along axes of alertness, contentedness and calmness (Norris, 1971). Participants completed this VAS three times during each session: immediately prior to drug administration (time 1), +90 min (immediately prior to cognitive testing, time 2) and +180 min (at the end of testing, time 3). Repeated measures ANOVAs were performed with Drug and Time as conditions. These revealed a main effect of Drug on Alertness (F(3,96) = 5.11, p = 2.53e-03) whereby Alertness during MPH administration was greater than PLA (t(32) = -3.14, p = 0.003), and not significant for ATM or CIT (p > 0.3). A main effect of Time on Alertness was present with decreasing Alertness across time (F(2,64) = 23.88, p = 1.79e-08). There was also a Time x Drug interaction (F(6,192) = 4.9, p = 1.08e-04), such that Alertness for MPH was not significantly different from PLA at Time 1 (t(32) = 0.28, p = 0.77), but was so at Time 2 (t(32) = -2.9, p = 0.006) and Time 3 (t(32) = -3.56, p = 0.001). There were no significant differences in Alertness for ATM and CIT across at any time. Thus, it appeared that MPH staved off the time-on-task decrement in alertness that is typically seen on cognitive tasks compared to the other drugs (Dockree et al., 2017). An effect of Time was also found for measures of Contentedness (F(2,64) = 21.15, p = 8.87e-08) and Calmness (F(2,64) = 4.04, p = 0.02), with ratings for both decreasing with time. There was however no interaction of Drug and Time for ratings of Contentedness or Calmness.

### Materials and task procedures

Participants performed a 2-stimulus oddball task (Figure 1). Visual stimuli were presented on a dark grey background and participants were instructed to fixate on the centre of the screen. They were also asked to restrict any eye movements throughout the task, e.g. large saccades or blinks. Every 2075 ms a stimulus appeared on the screen for 75 ms. Standard stimuli consisted of a 2 cm diameter purple circle which appeared on 80% of trials. Target stimuli were a slightly larger purple circle (4 cm diameter) which appeared on 20% of trials. Participants were asked to make a speeded response to target stimuli using the response box placed in their right hand. The stimulus array was pseudorandomly designed such that between 3 and 5 standard stimuli were presented after any target stimulus. Thus, the minimum interval between P3-eliciting events was 8300 ms. Participants performed 1 block of this task, comprising 42 target trials.

**Figure 1:**
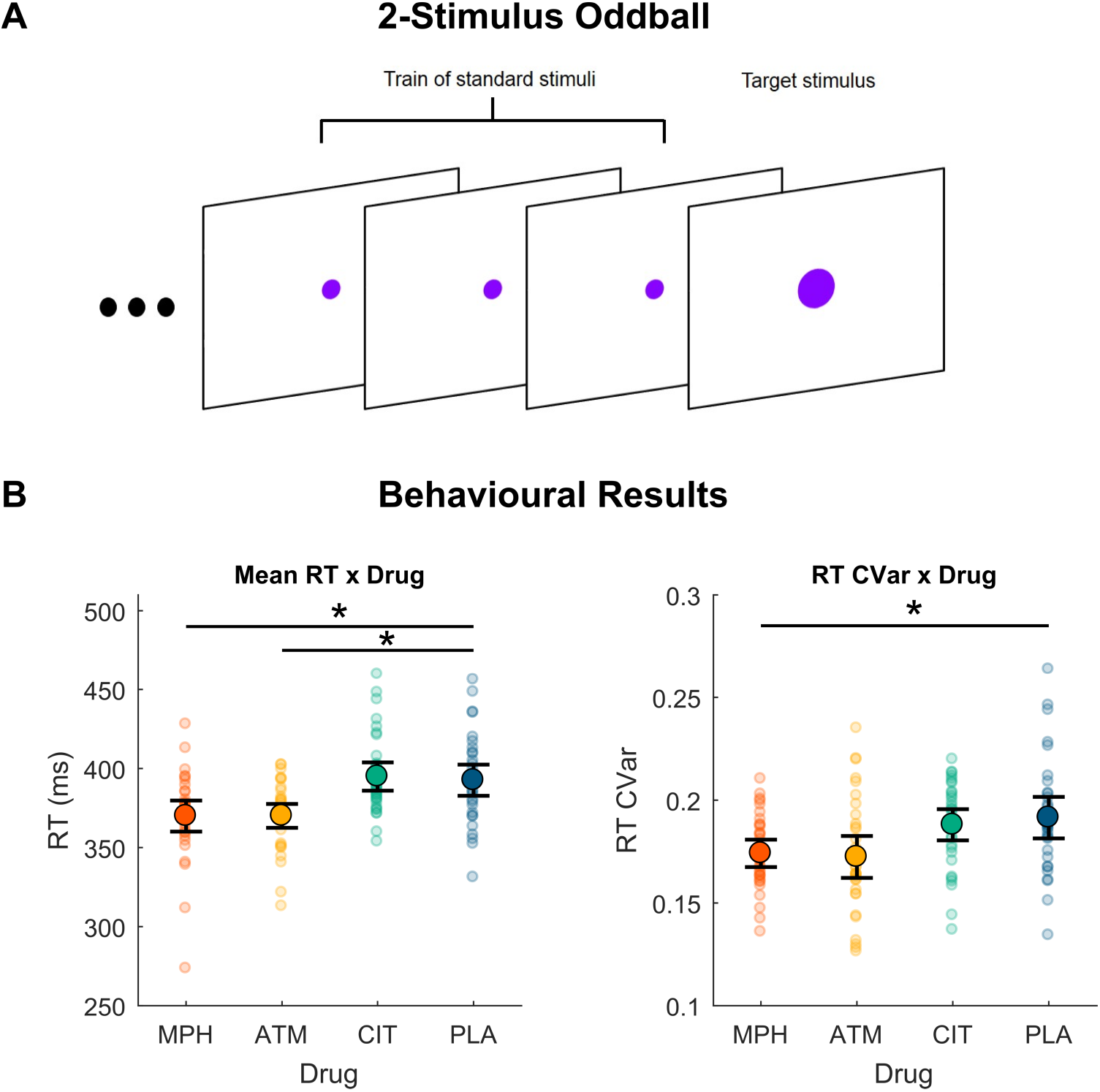
**A,** 2-stimulus oddball experiment. **B,** RT and RT coefficient of variability (RT CVar) x Drug. Error bars represent within-subjects 95% confidence intervals. Individual data points are mean centred for that individual by subtracting their overall value for the relevant measure (e.g. their mean RT), then adding the group mean. In this and the following figures, * denotes p<0.05, ** denotes p<0.01, *** denotes p<0.001.

### Experimental Design and Statistical Analysis

This experiment incorporated a within-subjects design, specifically using repeated-measures ANOVA and within-subjects t-tests, using the Benjamini & Hochberg procedure for controlling the false discovery rate (FDR) of a family of hypothesis tests (adjusted *p* values reported in text). Multilevel mediation analysis (Baron & Kenny, 1986) was performed in order to investigate the possible mediation of the drug effects on behaviour via our neural indices. The multilevel models involved in this analysis modelled between-subjects variability in our measures as random intercepts between subjects. Additional analyses are described in the Data Analysis, and the Results sections.

### EEG Acquisition and Preprocessing

EEG was recorded using an ActiveTwo BioSemi system of 64 scalp electrodes, sampled at 1,024 Hz. Data were preprocessed using MATLAB (MathWorks) and the EEGLAB plug-in (Delorme & Makeig, 2004). Data were resampled to 512 Hz for ease of processing, filtered with a 20-Hz low-pass filter and a 0.25-Hz high-pass filter and re-referenced offline to the average of all scalp electrodes. Epochs were then taken using a window from -125 ms before stimulus onset to 750 ms post-stimulus onset, and baseline corrected to the pre-stimulus period. To minimise interaction between overlapping ERP components, the data were subjected to Current Source Density transformation (Kayser & Tenke, 2006). A further analysis of pre-stimulus alpha used an epoch of 1000ms before target onset and alpha power (8 – 14Hz) was calculated using the Fast Fourier Transform of that timeframe. Trials were excluded from analyses if EEG from any channel relevant to the study (i.e. frontal channels and central parietal channels) exceeded +/- 100μV during the interval between 100 ms before target onset and the time of manual response. Critically, there were no significant differences in trial count across Drug conditions (F(3,96) = 0.65, p = 0.59; mean (SD) MPH - 36.7 (7.2), ATM - 35.9 (5.7), CIT - 35.6 (5.7), PLA - 38.0 (3.5)). Finally, in order to ensure single participants were not unduly influencing any analyses, participants were rejected from particular analyses if their data for those analyses were outside 3 standard deviations of the mean.

### Data Analysis

Behavioural performance measures testing the pharmacological manipulation were (a) reaction times (RTs) for correct detections, and (b) the coefficient of variability of RTs (RT Cvar), calculated per participant as the standard deviation of RT divided by mean RT (Bellgrove, Hester, & Garavan, 2004). Hit rate was at ceiling in this task (98.7% +/- 2%) and was not analysed further. There were no false alarms for standard stimuli.

We recorded the P3b ERP component in response to the targets as our neurophysiological measure of the evidence accumulation process. This allowed us to parse four separate aspects of decision formation which could theoretically explain the faster RTs: (i) P3b onset representing the start time of evidence accumulation; (ii) its build-up rate reflecting the rate of evidence accumulation; (iii) its amplitude at response execution representing the decision threshold; and (iv) its peak latency representing the time at which the threshold is reached (O’Connell et al., 2012; Twomey et al., 2015).

The P3b was measured from a parietal channel selected on a participant-by-participant basis, from 1 of 9 channels centred on CPz (the peak grand-average channel using the 10-20 coordinate system) as the channel with the greatest amplitude at manual response. Target-locked P3b epochs were -125 to 750 ms around target onset, and response-aligned P3b epochs were - 250 to 0ms around response.

We measured P3b onset via two methods. First, we calculated a running t-test against zero across time on the grand average P3b waveform (collapsed across subjects, and drug conditions) and identified the first time-point at which signal amplitude differed significantly from zero in a positive direction, at p<0.05. We then measured P3b onset as the amplitude from a 50ms window around this time-point. Second, we calculated P3b onset using a jackknife-based analysis (MILLER, PATTERSON, & ULRICH, 1998), which calculates the onset of a signal as the intersection of two lines, one fit to a defined pre-onset period (0 to 50ms post-stimulus onset) and forced to have a slope of zero, the other fit to a post-onset period (50ms to time-point of maximum amplitude of the signal). The 50ms cut-off between pre- and post-onset was chosen as the earliest possible onset of the P3b signal.

Build-up rate of evidence accumulation was calculated as the slope of a line fitted to the response-aligned P3b waveform from -150 to 0ms relative to response (Twomey et al., 2015). The 150ms window was chosen as one which captures the entire build-up of evidence up until manual response. P3b amplitude was measured at a participant-level in a 25ms window prior to response execution.

We measured the timing of stimulus-aligned P3b peak (latency) as the latency of peak amplitude in the trial-averaged stimulus-locked waveform, from grand average P3b onset (100ms) onwards. Averaged data were used in this case due to the difficulty of detecting peak latency in single trial data.

Visual Evoked Potential (VEP) components, the P1, N1, and P2 were analysed. Here, to maximise trial counts, both standard and target epochs were included. Again, relevant electrodes for each component were chosen on a per participant basis, the peak electrode chosen from a group of electrodes centred around the group average peak for that component.

To examine the effect of drug on the pre-stimulus power spectra, we performed a Fourier Transform on epochs from -1000ms to 0ms before stimulus onset. We hypothesised that prestimulus occipital alpha power (8 - 14 Hz) may be affected by drug (Dockree et al., 2017). We performed an additional exploratory analysis with Drug and Region (frontal, central, parietal, and occipital) as factors for the other frequency bands - theta (3 - 7 Hz), beta (15 - 30 Hz), and gamma (30 - 100 Hz).

## Results

Participants completed 4 sessions (MPH, ATM, CIT, or PLA) of the 2-stimulus oddball task, monitoring for a slightly larger diameter target circle pseudo-randomly appearing among a train of smaller standard circles (Figure 1A). Participants were faster to respond to the targets in the MPH and ATM conditions compared to the PLA condition (Figure 1B), whereas there was no significant difference in Reaction Times (RTs) between CIT and PLA (Main effect of Drug: F(3,96) = 7.14, p = 0.0002, partial *η*^2^=.18; MPH vs PLA: t(32) = -2.63, p = 0.02, 95% CI [-40.26, -5.09]; ATM vs PLA: t(32) = -3.03, p = 0.01, 95% CI [-37.76, -7.4]; CIT vs PLA: t(32) = 0.35, p = 0.72). Variability of RT (RT CVar) was also significantly lower in the MPH condition compared to PLA, with a similar trend in the ATM condition (Figure 1B): (Main effect of Drug: F(3,93) = 3.78, p = 0.013; MPH vs PLA: t(31) = -2.7, p = 0.03; ATM vs PLA: t(31) = -2.12, p = 0.06; CIT vs PLA: t(31) = -0.49, p = 0.63).

We found no difference in P3b onset across pharmacological conditions using either the amplitude of a 50ms window centered on the grand average peak (F(3,96) = 0.88, p = 0.46), nor the jackknife method (F(3,78) = 0.57, p = 0.64; Figure 2A(MILLER et al., 1998)).

**Figure 2:**
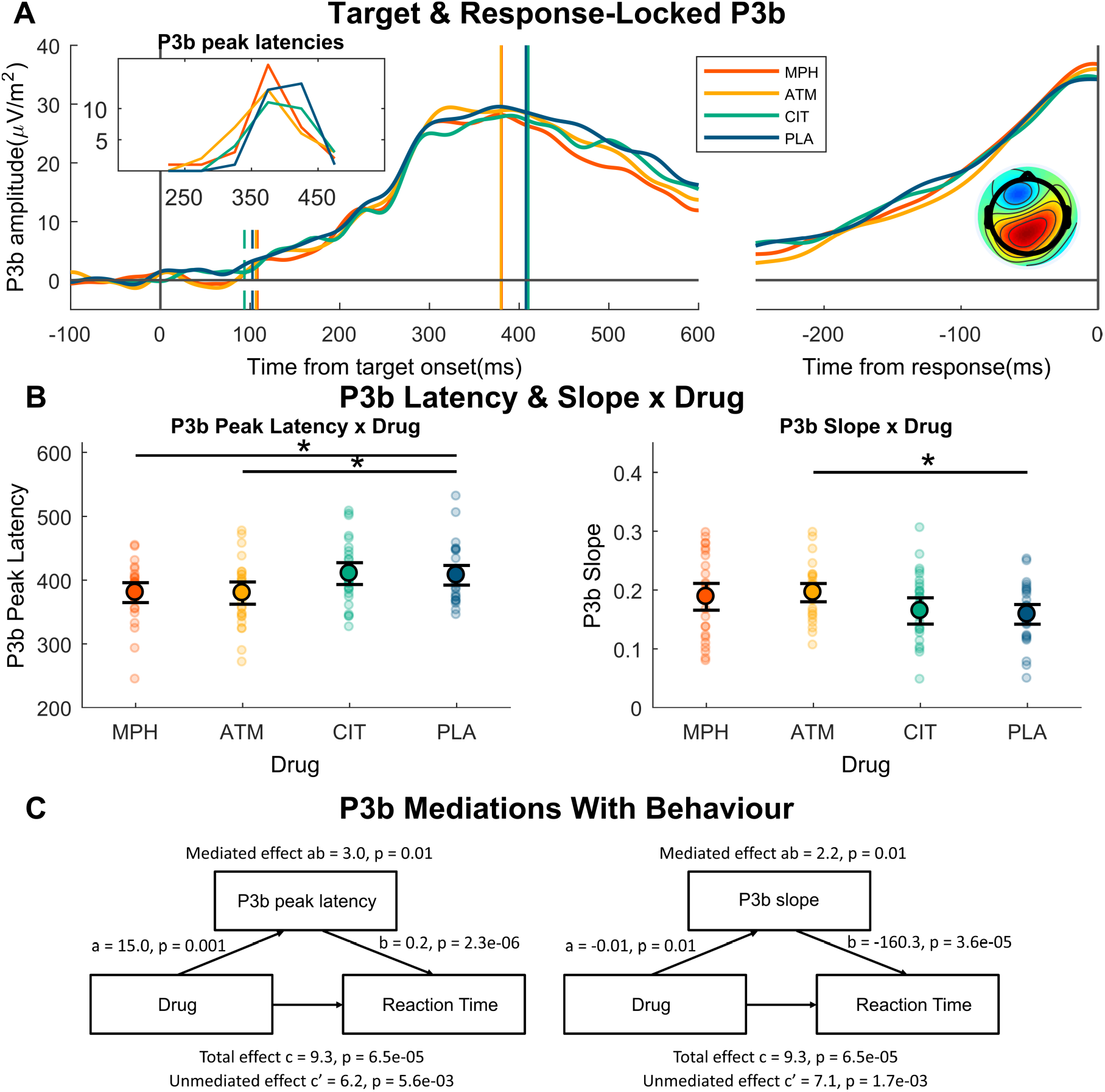
**A,** Target and response-locked P3b, plotted by drug. In the target-locked plot, the vertical dashed lines represent mean P3b onset latencies by drug. The vertical solid lines represent mean P3b peak latencies by drug. Note that MPH and ATM conditions have very similar peak latencies, making them difficult to discern here. **Inset left:** Histogram of P3b peak latencies showing the clear and reliable precedence of P3b latencies in MPH and ATM conditions compared to CIT and PLA. **Inset right:** Topoplot of the ERP at P3b peak latency. **B, Left,** P3b peak latencies x drug. **Right,** Response-locked P3b slopes x drug. **C,** Mediation parameters showing that the effect of drug on RT was partially mediated by **Left,** P3b peak latency and **Right,** P3b slope.

We found an effect of drug condition on P3b build-up rate (F(3,93) = 2.63, p = 0.05, partial *η*^2^=.08; Figure 2A, B), driven by a steeper P3b build-up in the ATM condition (t(31) = 2.83, p = 0.02, 95% CI [.01, .06]), and a similar trend in the MPH condition (t(31) = 1.9, p = 0.1, 95% CI [0, .06]). On the other hand, there was no effect of CIT on P3b build-up (t(31) = 0.4, p = 0.69). In order to verify that the effect of drug on behaviour was related to the effect of drug on P3b build-up, we performed a multilevel mediation analysis whereby we tested the hypothesis that the effect of drug on RT was mediated by the effect of drug on P3b build-up. We found that the effect of drug on RT was indeed partially mediated by the effect of drug on P3b build-up (Figure 2C).

A similar effect of drug condition was observed on the stimulus-locked peak latency of the P3b (F(3,90) = 3.22, p = 0.026, partial *η*^2^=.10; Figure 2A, B). This was driven by a shorter latency for both MPH and ATM (t(30) = -2.23, p = 0.05, 95% CI [-52.08, -53.05] and t(30) = -2.26, p = 0.05, 95% CI [-53.05, -2.65], compared to PLA respectively), whereas CIT was not significantly different from PLA (t(30) = 0.19, p = 0.85). Similar to the analysis of P3b build-up, we performed a multilevel mediation analysis, investigating the mediation of drug effect on RT via P3b peak latency. Again, we found a partial mediation effect whereby P3b peak latency mediated the effect of drug on RT (Figure 2C).

Next, we examined the possibility that drug could impact the speed of decision-making via a lowering or raising of the bound of evidence accumulation, here represented by the peak amplitude of the response-locked P3b (Twomey et al., 2015). There was no significant effect of drug condition on response-locked P3b peak amplitudes (F(3, 93) = 0.56, p = 0.64).

Finally, we performed an analysis to explore any possible differences across drug condition in the relative timing of each participant’s response-locked P3b peak compared to their average RT. In behavioural models of decision-making, this is reflected in a change in the non-decision time parameter along with the time between stimulus-onset and evidence accumulation onset (Smith and Ratcliff, 2004; Ratcliff and McKoon, 2008). We found no difference in the relative timing of P3b peak compared to RT across drug conditions (F(3,96) = 0.34, p = 0.79).

Analysis of the VEPs revealed no no significant effect of drug condition on P1 or N1 amplitude. There was however a significant drug effect on P2 amplitude (F(3, 96) = 5.4, p =0.001, partial *η*^2^=.14), driven by a greater P2 amplitude in the MPH condition compared to PLA (t(32) = 2.68, p = 0.03, 95% CI [.93 6.83]). There was, however, no significant mediation of the drug effect on RT by P2 amplitude.

**Figure 3:**
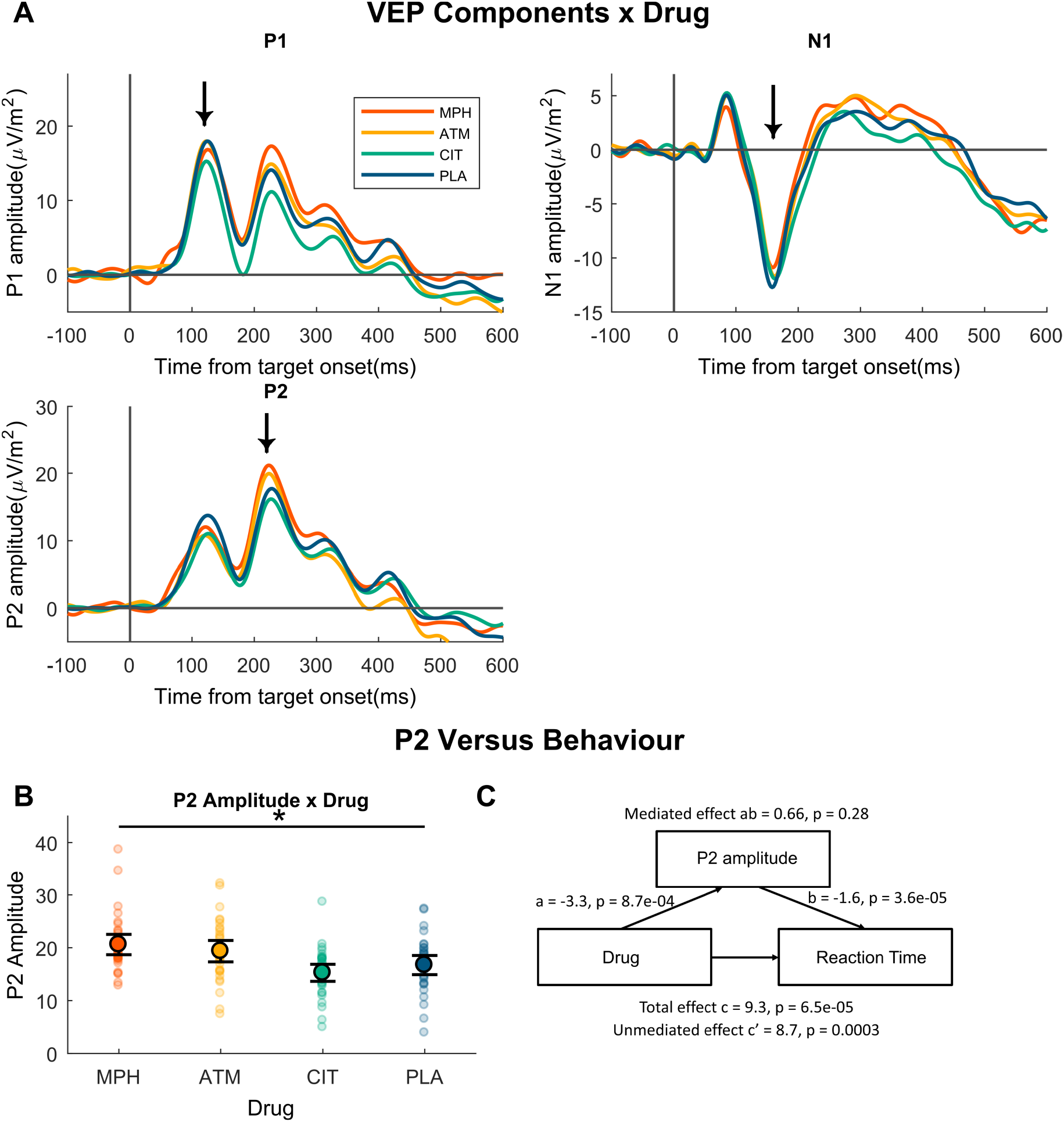
**A,** P1, N1, and P2 components by drug condition. **B,** P2 peak amplitudes x drug. **Right,** Mediation parameters showing that the effect of drug on RT was not mediated by P2 peak amplitudes.

We found no differences in pre-stimulus posterior alpha power (7 - 14 Hz) between any of the drug conditions (F(3,96) = 0.82, p = 0.49, Figure 4B). We found a reliable decrease in theta power across the scalp in the MPH and ATM, but not CIT conditions (Main effect of drug: F(3,96) = 5.64, p = 0.001). Given the inverse relationship between theta power and arousal levels (Klimesch, 1999), it is possible that this effect was due to increased arousal during the MPH and ATM conditions. Second, there was a broadband increase in beta/gamma power during the CIT condition compared to the other conditions, focussed on fronto-temporal areas (Interaction effect of drug x ROI: F(9,288) = 5.66, p = 3.2e-07). The broadband nature of this effect would suggest that it could be related to muscular high-frequency artifacts (e.g. jaw-clenching).

**Figure 4:**
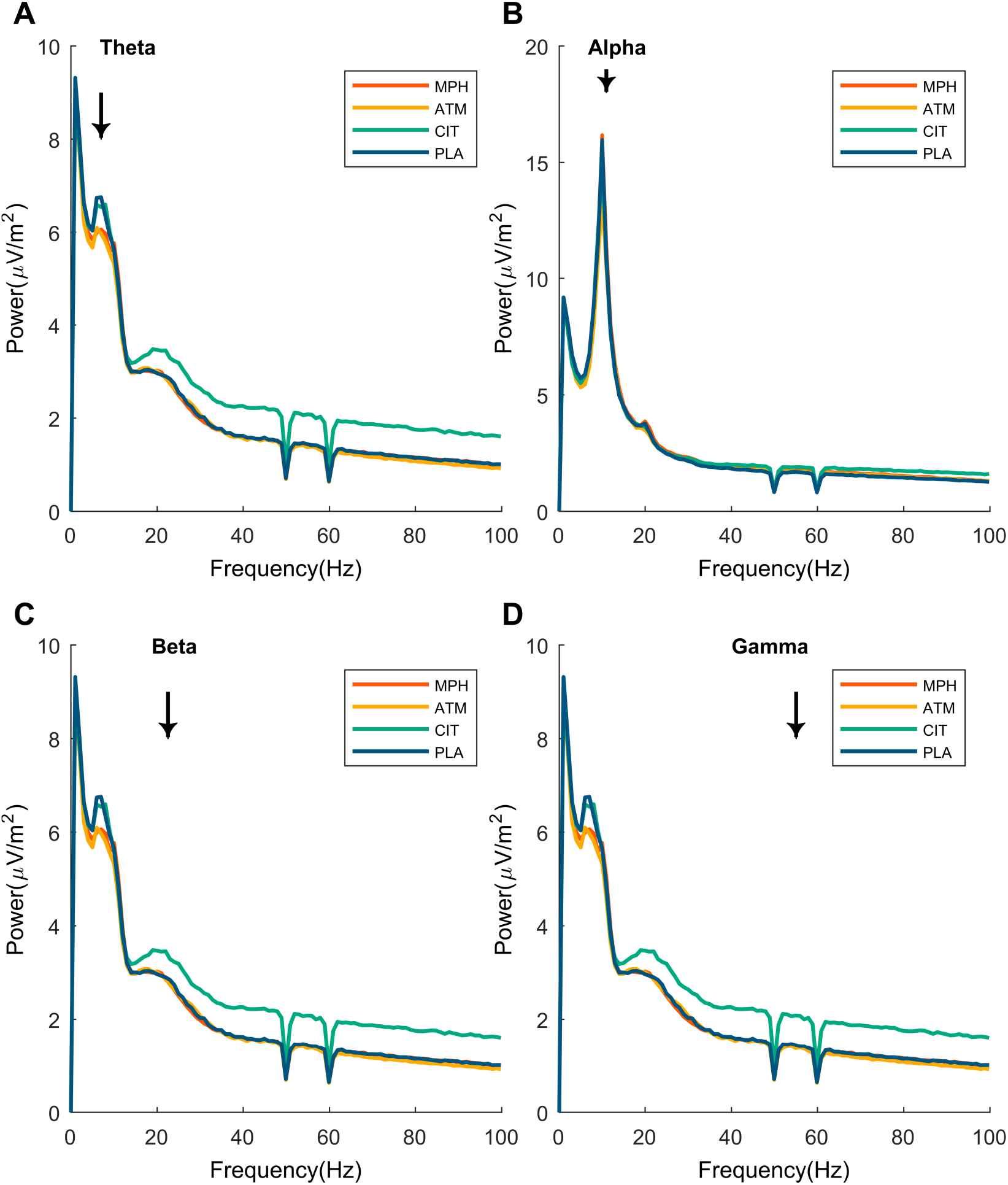
Pre-stimulus power spectra by drug condition **A,** Theta, **B,** Alpha, **C,** Beta, **D,** Gamma. Each spectrum is plotted from the region of greatest difference between drug conditions for a particular frequency band. Notch filters at 50 and 60Hz are present.

## Discussion

The present results provide direct neurophysiological evidence of catecholaminergic modulation of the rate of evidence accumulation during perceptual decision-making. Here, we employed a randomised, double-blinded, placebo-controlled, crossover design combined with the classic 2-stimulus oddball task. We found that the introduction of MPH and ATM, but not CIT, speeded RTs to the oddball targets compared to a placebo condition. This behavioural facilitation was accompanied specifically by faster build-up rates and earlier P3b peak latencies in the MPH and ATM conditions, with no change in either the onset or peak amplitude of the P3b. This points to an effect of noradrenaline, and possibly dopamine, on the rate of evidence accumulation towards target detection in the human brain.

Catecholaminergic speeding of target detection is a well-established finding across a variety of tasks (BRUMAGHIM & KLORMAN, 1998; Linssen et al., 2014; Naylor et al., n.d.). Recently, Fosco and colleagues(Fosco et al., 2016) used a drift diffusion model (DDM) in ADHD to show that MPH increased the drift rate and non-decision time, while reducing boundary separation. The present study provides neurophysiological support for one aspect of their behavioural analyses-steeper build-up rate of the P3b by MPH, representing a faster rate of evidence accumulation towards response (O’Connell et al., 2012; Twomey et al., 2015). Fosco et al. found that MPH increased the non-decision time (the amount of RT accounted for by decision-independent processes such as sensory encoding and motor execution(Ratcliff & McKoon, 2008; Smith & Ratcliff, 2004)) component of their DDM. Given the beneficial effects of MPH on behaviour, this observation was interpreted as counterintuitive. We did not find any effect of MPH or ATM on P3b onset or on the relative timing of P3b peak compared to RT; either of these effects would have been consistent with a modulation of non-decision time processes. Furthermore, Fosco et al. found decreased boundary separation in their model, an effect which could manifest itself in our task as a decrease in CPP amplitude. We propose this discrepancy is due to task differences. In contrast to our simple detection task, Fosco et al. used a two-choice discrimination task, which could have engendered a strategic need for boundary separation in order to discriminate accurately. Finally, the findings of Fosco et al. derive in part from individuals with ADHD and thus may not be directly comparable to the current study with neurologically healthy participants

MPH and ATM increase levels of both dopamine and noradrenaline to varying extents in different brain regions, and it is difficult to disentangle their dopaminergic and noradrenergic effects (Berridge et al., 2006; Bymaster, 2002; Han & Gu, 2006). Our results are nevertheless in agreement with recent work showing an increase in the precision, or signal-to-noise ratio, of cortical representations due to catecholaminergic intervention(Warren et al., 2016). First, we show a catecholamine-related increase in the rate of evidence accumulation, which could theoretically occur either due to an increase in neural noise or more precise representation of the target. Second, we also show a concurrent decrease in RT variability which would imply the latter scenario of increased signal-to-noise ratio of neural representation of the target, as opposed to increased neural noise which would result in greater RT variability.

The lack of catecholaminergic effect on the early components of the VEP, i.e. P1 and N1, points to a noradrenergic influence on relatively later stages of information processing. The adaptive gain theory of the Locus Coeruleus-Noradrenaline system (LC-NE), attributes the influence of phasic Noradrenaline-related LC responses to the translation of post-threshold activity from the decision layer to the response layer (Aston-Jones & Cohen, 2005; Nieuwenhuis et al., n.d.). In our data, this would appear as a decrease between P3b peak latency and RT, which was not observed. Specifically, our results suggest an effect at the timing of the decision layer. This could occur as neural gain between the sensory and decision layers for example, where decision-relevant neurons are more sensitive to input from sensory regions (Warren et al., 2016).

The present findings are consistent with those which have previously examined catecholaminergic effects on the P3b. Drugs which increase the amount of noradrenaline in the system have been shown to result in a faster P3b peak latency (BRUMAGHIM & KLORMAN, 1998; Dockree et al., 2017; Naylor et al., n.d.), whereas drugs such as clonidine which decrease noradrenaline signalling have been shown to result in slower peak latency (HALLIDAY et al., 1994; Swick, Pineda, & Foote, 1994). Other studies have shown an increase in P3b peak amplitude in response to MPH, which we do not confirm here (BRUMAGHIM & KLORMAN, 1998; COOPER et al., 2005; Dockree et al., 2017). The absence of a drug effect on P3b amplitude in the present study could arise from a ceiling effect for performance (Hit rate 98%) whereby the evidence accumulation process reliably reaches a bound regardless of pharmacological manipulation because of the ease of our task. Although our results generally agree with past studies of catecholamine modulation of target detection using EEG, our dissection of the P3b into its distinct parameters (onset, build-up rate, peak latency and amplitude) provides unique mechanistic insight into the impact of catecholamines on the neural decision process. Such a framework may be profitably applied in future pharmacological studies to inform the pharmacology of discrete processing stages underpinning human choice behaviour.

Although MPH and ATM are the most universally prescribed psychostimulants for the treatment of ADHD (Garnock-Jones & Keating, 2009; Storebø et al., 2012), we lack a clear understanding of the neurophysiological bases of their ability to enhance behaviour. Such insights are critical for the identification of robust biomarkers of drug response that may ultimately facilitate personalized approaches to treatment in disorders such as ADHD. Our recent work has demonstrated the effect of MPH on attentional engagement which was linked to a suppression of alpha-band activity as well as a reduction in its trial-to-trial variability (Dockree et al., 2017). Interestingly, we observed no such drug effects here despite the fact that our data were collected from the same participants and in the same testing session. In our view the differences in results arise from differences in the tasks employed. The oddball task employed here is relatively easy, with performance close to ceiling, whereas the task employed by Dockree et al was a very demanding sustained attention task designed to engender lapses of attention (hit rate was approx. 60 - 75%). Thus, it is plausible that the oddball task did not engender behaviourally relevant fluctuations in attentional engagement, thus leaving little scope for modulation by drug. This highlights the potential for medications such as MPH to exert task-dependent effects on neural activity and the utility of neurophysiology for dissecting these contributions.

## Acknowledgements

This work was supported by Grant No. 569532 from the National Health and Medical Research Council of Australia (to MAB) and Young Investigator Grant No. 22457 from the Brain and Behavior Research Foundation (to RGO). MAB is supported by a Future Fellowship No. FT130101488 from the Australian Research Council. This project was also supported by Marie Curie International Research Staff Exchange Scheme No. 612681 under the European Commission FP7 (to RGO and MAB). This study was registered on the Australian and New Zealand Clinical Trials Registry: The Effect of Methylphenidate, Atomoxetine and Citalopram Versus Placebo on Behavioural and Physiological Indices of Executive Control in Healthy Individuals; http://www.anzctr.org.au/ACTRN12609000625279.aspx; ACTRN12609000625279.

